# A fully automatic method yielding initial models from high-resolution electron cryo-microscopy maps

**DOI:** 10.1101/267138

**Authors:** Thomas C. Terwilliger, Paul D. Adams, Pavel V. Afonine, Oleg V. Sobolev

## Abstract

A fully automated procedure for optimization and interpretation of reconstructions from cryo-EM is developed and applied to 476 datasets with resolution of 4.5 Å or better, including reconstructions of 47 ribosomes and 32 other protein-RNA complexes. The median fraction of residues in the deposited structures reproduced automatically was 71% for reconstructions determined at resolutions of 3 Å or better and 47% for those at lower resolution.

---

Electron cryo-microscopy (cryo-EM) has recently come of age for determining three-dimensional structures of large biological complexes^1,2^. Improvements in data collection and image reconstruction methods have made it possible to obtain 3D images of macromolecules at resolutions where structural details such as locations of side chains can be readily visualized. A key limiting factor in structure determination by cryo-EM is now the effort required for interpretation of these reconstructed images in terms of an atomic model. As cryo-EM structures often have many thousands of protein or nucleic acid residues, this manual interpretation is very time-consuming. Currently, reconstructions are typically interpreted by identifying regions of the map containing individual components, adjusting the sharpening of the map, and fitting previously-determined structures^3^ or building a model *de novo* into the image using graphical tools^4^ or automatic interpretation of the map^5–8^.

Algorithms for *de novo* model-building using cryo-EM maps have been developed recently^5–9^ and are capable of building very complete models in some cases. However, they do not integrate model-building with the other major steps in map interpretation. None, for example, can carry out map segmentation, automatically sharpen a map, identify what parts of the map correspond to protein, RNA or DNA when more than one type of chain is present in the same map, or take reconstruction symmetry into account. Consequently they are not easy to apply on a global scale, and each has been demonstrated with only a few cryo-EM reconstructions. In the recent cryo-EM Model Challenge^10^, for example, there were no submissions for automatic model-building (other than those using the methods described here) for 5 of the 12 cryo-EM maps in the challenge, including the two ribosome reconstructions (TT and PDA, unpublished).

Our fully integrated and automated procedure carries out map interpretation without user intervention and requires as inputs only a cryo-EM map, the nominal resolution of the reconstruction, the sequences of the molecules present, and any symmetry used in the reconstruction. Map interpretation begins with automatic image sharpening using an algorithm we have recently described^11^ that maximizes connectivity and detail in the map. The unique parts of the structure in the sharpened map are then identified with a new automatic segmentation algorithm that takes into account any reconstruction symmetry (See Online Methods for details of all new methods). For each part of the structure and for each type of macromolecule present, atomic models are generated using several independent methods. Our feature-based searches for protein secondary structure^12^ and template-based protein main-chain building^13^ have been described previously. Here we describe methods for automatic nucleic acid model-building, we introduce a new method for protein model-building that uses our chain-tracing algorithm^14^ followed by extensive real-space refinement^15^ using restraints based on automatically-identified secondary structure elements, and we introduce new methods for recombination among all models for all chain types present, based on fit to the reconstruction. Finally our automated procedure carries out assembly and refinement of the entire structure including any symmetry that is present.

Fig. 1A illustrates part of a deposited cryo-EM reconstruction for lactate dehydrogenase (EMDB^16^ entry 8191) obtained at a resolution of 2.8 Å, along with an atomic model (PDB^17^ entry 5k0z^18^) representing the authors’ interpretation of the reconstructed map. This reconstruction has D2 symmetry with 4 copies of one unique chain. In the process of fitting an X-ray structure into the map to obtain this model, the authors used sharpened versions of the map to enhance high-resolution features and allow more accurate model-building^18^. Our automated procedure optimizes map sharpening and yields an atomic model as an interpretation of that map (Fig. 1B). In the sharpened map in Fig. 1B, features such as side-chains can be clearly visualized without the need for a researcher to manually adjust the sharpening of the map, though our automated interpretation in Fig. 1B does not detect these side-chains in this case.

**Fig. 1.**
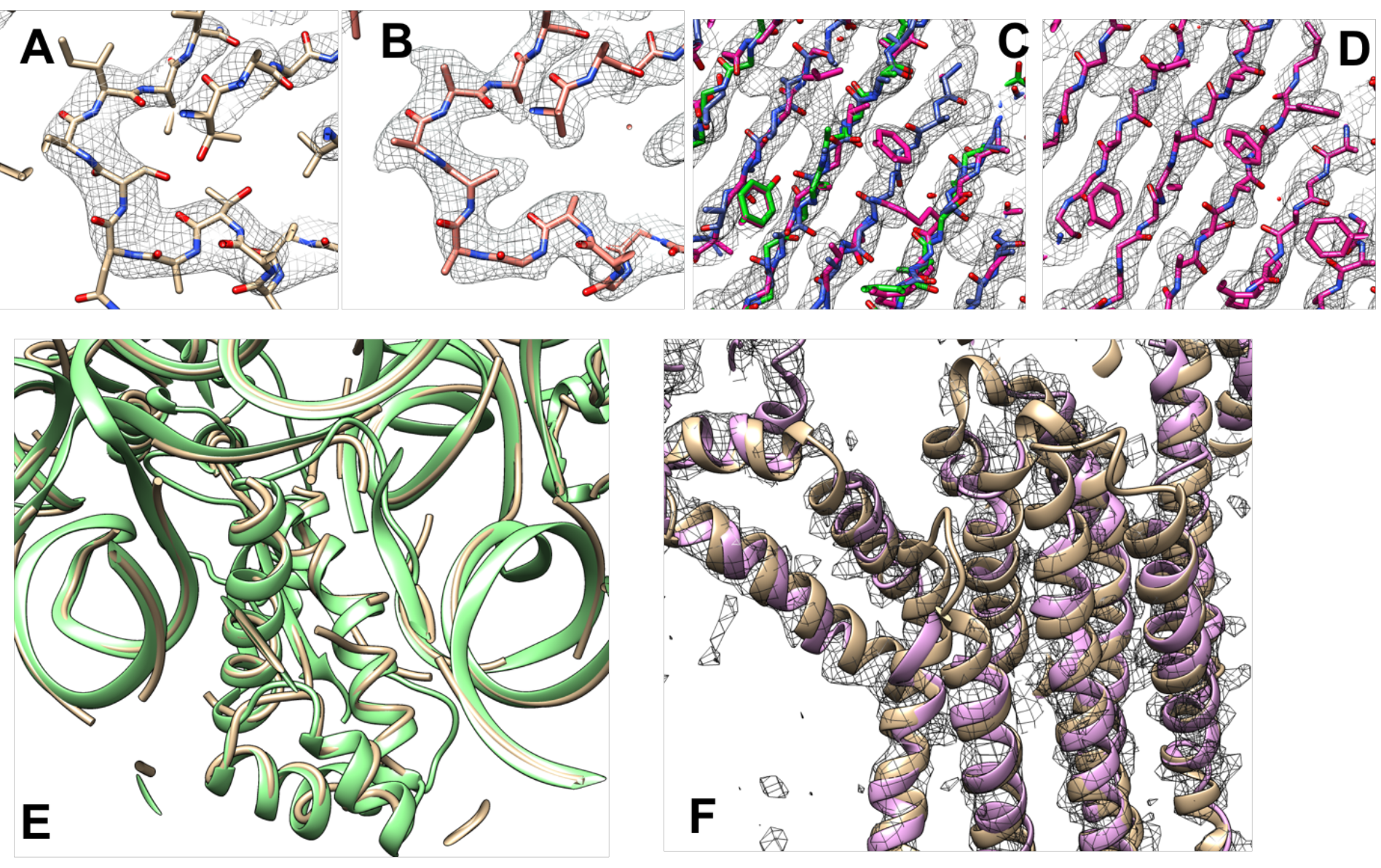
Automated interpretation of cryo-EM maps. A. Section through deposited cryo-EM reconstruction and deposited interpretation for lactate dehydrogenase^18^ (EMDB entry 8191). B. Automatically sharpened version of map in A with automatically generated model (PDB entry 5k0z). C. Automatically sharpened version of map for EMDB entry 6630 (glutamate dehydrogenase^19^) with three independent automatically-generated interpretations (using chain-tracing, green model, feature-based helix and strand identification, blue model, and pattern-based secondary structure identification with fragment-based extension, magenta model). D. Composite model derived from the three models in C. E. Automatically interpreted reconstruction of the *Mycobacterium smegmatis ribosome* (yellow tubes) compared with deposited model (green ribbons). F. Automatically generated model of the ERAD channel Hrd1 (purple ribbons) compared with deposited model (brown ribbons). Note in each case that most portions of the automatically generated model match the deposited model and a few portions do not. See text for details. Graphics created with Chimera^20^.

Fig. 1C illustrates the procedure in more detail for the deposited cryo-EM reconstruction for glutamate dehydrogenase (EMDB^16^ entry 6630) obtained^19^ at a resolution of 3.3 Å. Using the automatically sharpened map, three models were created, each using a different algorithm so as to minimize correlated errors. These three models are superimposed on the map in Fig. 1C (after applying reconstruction symmetry). The best-fitting parts of each model were combined and extended to fill gaps and the resulting composite model was refined to yield an interpretation of the reconstruction (Fig. 1D). For this glutamate dehydrogenase structure, 66% of the residues in the deposited model were reproduced by the automatically-generated model, with a root-mean-square (rms) coordinate difference of 0.8 Å for C_α_ atoms.

We used a similar procedure to automatically generate interpretations of cryo-EM reconstructions containing complexes of RNA and protein. In addition to generating multiple models interpreting each part of a map as protein, further models were generated interpreting them as RNA, and the best-fitting parts of each model were combined to represent the protein-RNA complex. Fig. 1E shows such an automatically interpreted reconstruction of the *Mycobacterium smegmatis ribosome*^21^ (EMDB entry 6789) analyzed at a resolution of 3.1 Å and compared with the deposited model. The automatically generated model represents 60% of the RNA and 48% of the protein in the deposited model (PDB entry 5xym), with rms coordinate differences of 0.7 Å for P atoms and 1.0 Å for C_α_ atoms.

At lower resolution, a smaller fraction of the structures are typically reproduced by our methods, but secondary structural elements such as protein or RNA helices are often still identifiable. Fig. 1F illustrates the automatically sharpened map and compares the automatic interpretation for the protein-conducting ERAD channel Hrd1^22^ (EMDB entry 8637) with the deposited model. This reconstruction was previously interpreted using a combination of fitting helices into density, manual model-building and Rosetta modeling with distance restraints from evolutionary analysis22 (PDB entry 5v6p). Our automatic approaches reproduce the positions of 81% of the residues in the model with rms coordinate differences for matching C_α_ atoms of 1.3Å.

We evaluated the overall effectiveness of our procedure by applying it to all 476 high-resolution cryo-EM reconstructions in the EMDB database that we could extract with simple tools and match to an entry in the PDB. The information supplied to our algorithm for structural interpretation consisted of the deposited reconstruction, the sequences of the molecules present, and the resolution and symmetry of the reconstruction. We used reported resolutions of the reconstructions (ranging from 1.8 Å to 4.5 Å) and deduced the symmetry used in reconstruction from metadata present in the databases and from the symmetry in the deposited models. We use the deposited models as our best available representation of the correct structure in each case. Fig. 2A shows the fraction of residues in each deposited model reproduced by our analysis as a function of resolution of the reconstructions. Fig. 2B shows the rms coordinate difference between the generated model and deposited model corresponding to each map. Fig. 2C shows the fraction of these matching residues that also shared the residue type of the corresponding residues in the deposited structure. For chains in maps reconstructed at resolutions of 3 Å or better, the median fraction of protein and RNA residues in the deposited models reproduced by our approach was 71% and 45% respectively. At lower resolutions, 47% of protein residues and 34% of RNA residues were reproduced. The median rms coordinate differences for matching C_α_ atoms in protein chains and for matching P atoms in RNA chains were each about 1/3 the resolution of the reconstructions (Fig. 2B). The median fraction of the sequence of the deposited structure that could be reproduced was 28% for protein chains at higher resolution and 9% at lower resolutions (where random is about 5%). For RNA chains the sequence match was 49% at higher resolution and 42% at lower resolutions (random is about 25%). This higher sequence match for RNA chains is likely made possible both by the smaller number of possibilities (4 vs 20) and by our approach of considering base-paired residues together, effectively doubling the amount of density information available for identification of each base.

**Fig. 2.**
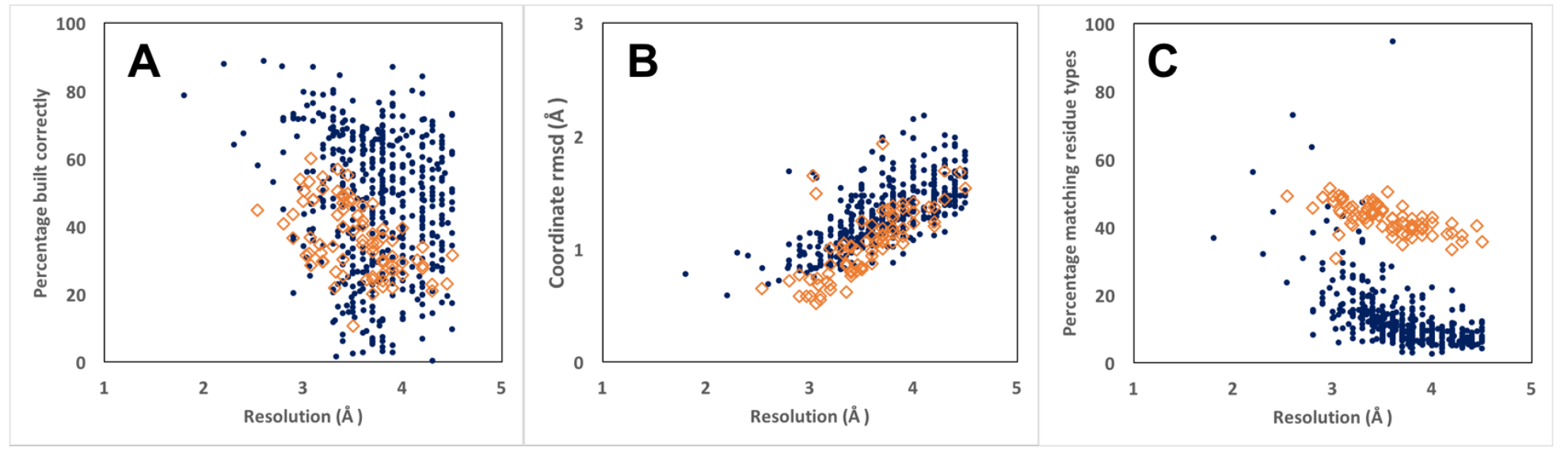
Residues in deposited models reproduced by automated analysis of cryo-EM reconstructions. Protein chains are indicated by blue dots and RNA chains are indicated by open brown diamonds. Residues are considered matched if C_α_ atom or P atom coordinates match within 3 Å (see Methods). Rms coordinate differences are assessed forall matched residues. A. Fraction of residues in deposited structure reproduced by automated analysis. B. Rms coordinate differences between matched residues in deposited structure and automated analysis. C. Fraction of residues in automated analysis matching a residue in the deposited model that also share the residue type of the matching residue in the deposited structure.

With our automated procedure, essentially any high-resolution reconstruction with suitable meta-data describing the reconstruction, its resolution and reconstruction symmetry can be interpreted and a first atomic model generated without any manual intervention. Although the models produced are not complete at this stage, we anticipate that combining the integration available in approaches such as ours with the greater completeness achievable in some cases with others’ algorithms^5–9^ may lead to both a high degree of automation and high model completeness.

Full automation makes analyses possible that would be challenging to carry out with tools that require manual input. One such important class of analyses is error estimation. With automated interpretation of a map, error estimates can be obtained by repeating the interpretation using different algorithms or different random seeds in appropriate stages of analysis^23,24^. The variation among models all generated using the same data and all in similar agreement with the data and prior knowledge yields an estimate of the errors in these models. If the errors in the models are independent, this variation can be an accurate estimate of errors, while if the models have correlated errors, it is rather a lower bound on the errors in these models^23^.

We illustrate two applications of repetitive map interpretation to error estimation. Above in Fig. 1C, three models were generated based on a sharpened cryo-EM map using different algorithms. The differences between overlapping parts of these models can be used as an estimate of the uncertainties in the models^23^. We first carried out a simulation to verify that this procedure would be expected to work in an ideal case. We generated maps with varying amplitude and phase errors based on chain A of PDB entry 5k0z^18^ at a resolution of 2.8 Å, and we then analyzed each map with our standard procedure. For each map, we calculated the true error in the models we generated based on their coordinate rmsd to the known true structure. We also calculated the coordinate rmsd between the two independent automatically-generated models, which would be expected to be about √2 times their individual errors if they are independent. In Fig. 3A we compare these uncertainty estimates with the actual coordinate differences, including only cases where at least 50% of the known structure was reproduced so as not to include very poor models. We find that they are similar to expected values (the line has a slope of √2), though with a small systematic difference that is consistent with a small correlation of errors in the automatically-generated models. In Fig. 3B we apply the same analysis to the models generated from data in the EMDB and compared with deposited models. Fig 3B shows that this relationship is very similar to the one shown in Fig. 3B, except that the slope is slightly different. The slope indicated by the line in Fig. 3B corresponds to that expected if the deposited models each had about half the rms error of the automatically-generated models. This analysis suggests that internal consistency of independently-generated models may be useful in creating estimates of model error. Related estimates could be obtained by interpretation of independent half-datasets as suggested recently^24^.

**Figure 3.**
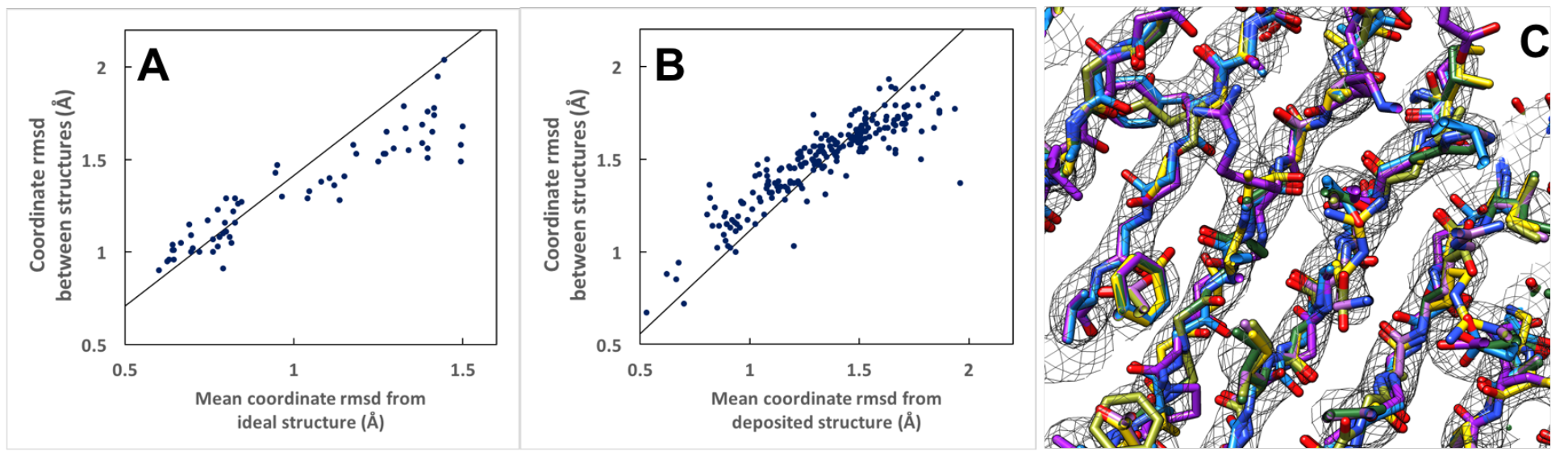
Applications of automated map interpretation to error analysis. A. Simulation with known true structure, maps with simulated errors, and automatic interpretations of maps with errors. Comparison of rms coordinate differences between automatically-generated models and the known true structure (average of two values) with coordinate differences between the two automatically-generated models. The slope of the line (√2) represents the ideal slope if errors were random and equal for the two automatically-generated models. Models were generated by chain tracing followed by iterative coordinate randomization and refinement with automatically-determined secondary structure restraints, and by pattern recognition of secondary structure elements followed by iterative extension with short peptide libraries. B. Analysis of data from analysis of maps from the EMDB as in A, except the ordinate values correspond to the mean difference from the deposited structure. The line shown corresponds to the slope expected (approximately 1.1) if errors in the deposited model were half the size of those in the automatically-generated models. C. Six models resulted from repetitive interpretation of sharpened map in Fig. 1C with varying random seeds. See text for details.

Fig. 3C illustrates how repetitive map interpretation can be used to estimate local uncertainties in automatically-generated models. The sharpened map in Fig. 1C was interpreted six separate times with *phenix.map_to_model*, each time with a different random seed for all steps such as refinement and fragment extension where randomization is used. The six models generated in this way are superimposed in Fig 3C, and it can be readily appreciated that the model shown in Fig. 1C was not the only possible interpretation of this map. This analysis gives an idea of the uncertainty in the model and potentially in each part of the model, with caveats as discussed previously in the context of crystallographic models^23^.

In addition to providing mechanisms for error estimation in atomic models representing cryo-EM reconstructions, the full automation of map interpretation provides a path towards the vision of continuous re-interpretation of deposited cryo-EM reconstructions and improvement of the models that represent them^25^. In the crystallographic field, deposited structures are already being continuously improved with database-wide re-refinement procedures^26^. As improved procedures are developed for interpretation of cryo-EM maps, the new procedures can be incorporated into automated frameworks for re-interpretation of all existing data, yielding ever-improving representations of these structures. This re-interpretation could eventually be extended to begin with the original images obtained from cryo-EM, as is beginning to be done with X-ray data^27^.

## Methods

### Automatic map interpretation

The steps required for map interpretation are all carried out automatically by the *Phenix*^28^ tool *phenix.map_to_model*. Here we describe the steps in detail, focusing on the methods that have not been previously reported, along with *Phenix* tools that can be used to carry out individual steps.

### Map optimization

Maps are automatically sharpened (or blurred) with *phenix.auto_sharpen*^11^, which maximizes the level of detail in the map and the connectivity of the map by optimizing an overall sharpening factor^29^ *B*_*sharpen*_ applied to Fourier coefficients representing the map up to the effective resolution of the map, beyond which a blurring exponential factor *B*_*blur*_ with a value of 200 Å^2^ is applied.

Map segmentation is carried out by identifying all regions of density above an automatically-determined threshold, choosing a unique set of density regions that maximizes connectivity and compactness, taking into account the symmetry that is present. In order to account for variable density levels in different parts of the map, the process is repeated with a new automatically-determined threshold after removing all the density that has been accounted for in the first iteration.

Contiguous regions above a threshold in a map are identified using a region-finding algorithm. This algorithm chooses all the grid points in a map that are above a given threshold. Then it groups these grid points into regions in which every point in a region is above the threshold and is connected to every other point in that region through adjacent grid points that are also above the threshold.

The choice of threshold for defining regions of density is a critical parameter in segmentation^30^. We set this value automatically by finding the threshold that optimizes a target function that is based on three factors. One factor targets a specific volume of the map above the threshold. A second factor targets the expectation that if n-fold symmetry is present, then groups of n regions should have approximately equal volumes. A third factor is targets regions of density of a specific size. The desired volume above the threshold is chosen based on the molecular volumes of the molecules expected to be present in the structure and the assumption that a fraction *f*, (typically 0.2) of the volume inside a molecule will have high density (the parts very near atoms) and that only these high-density locations should be above the threshold. The desired size of individual regions of density is set to be about the size occupied by 50 residues of the macromolecule, chosen because this is a suitable size for model-building of one or a few segments of a macromolecule. The exact size of regions is not crucial; it is just difficult to build a low-resolution structure looking at one residue at a time and unnecessary to build one looking at 1000 residues at a time.

The details of setting this threshold depend on *n*, the number of symmetry copies in the reconstruction, *n*_*res*_ the total number of residues in the reconstruction, the total volume of the reconstruction *v*_*total*_, the volume occupied by the macromolecule, *v*_*protein*_, a target fraction f of grid points inside the macromolecular regions desired to be above the threshold, and a target number of residues in each region of *r.* Prior to segmentation, the map is normalized, taking into account the fraction *v*_*protein*_ /*v*_*total*_ of volume of the map that contains macromolecule. The map is first scaled such that its mean value is zero and variance is unity. Then a scale factor of

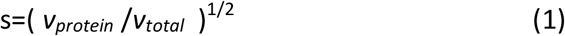
is applied. This scaling has the property that if the density for the molecule has uniform variance everywhere inside the molecule, removing part of the molecule from the map would lead to a scaled map that is unchanged for the remainder of the molecule.

Now the desired total volume *v*_*target*_ corresponding to high density within the macromolecule is given by the product of the total volume within the macromolecule, *v*_*total*_, and the desired fraction of grid points within the macromolecule that are to be above the threshold, *f*,

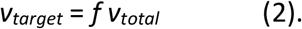

The number of desired regions *m*_*target*_ is given by the number of residues in the macromolecule, divided by the desired residues in each region,

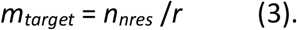

The desired volume per region *v*_*region_target*_ is the ratio of the total target volume to the total number of regions,

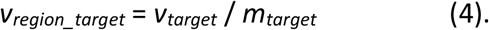

The desired volume ratio of the n’th region, *v*_*n*_, to that of the first, *v*_*1*_, is unity, and the value of this ratio is,

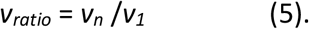

For a specific threshold, the volumes of regions above the threshold and the median volume *v*_*median*_ of the first *m*_*target*_ of these regions, after sorting from largest to smallest, are noted.

The desired median volume *v*_*median*_ is *v*_*region_target*_. We use the target function,

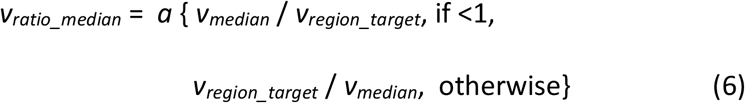

to express this, so that a high value of *v*_*ratio_median*_ is always preferred. If all regions are about equal in size then this volume ratio *v*_*ratio_median*_ is not informative. The weight on the volume ratio is therefore scaled, increasing with variation in the size of regions, using the formula,

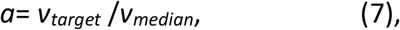

where *a* is expected to increase from a value of 1 if all regions are the same size, to larger values if regions are of different sizes, as the largest regions will have more than median volumes.

The desired volume of the largest region, v1, is also vregion_target. The target function,

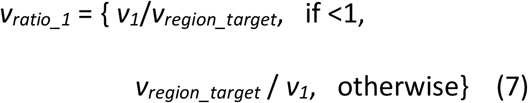

expresses this.

Finally, empirically we find that a threshold *t* on the order of unity is typically optimal (after scaling of the map as described above). We express this with a final ratio,

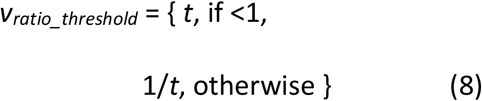

where larger is again desired.

The total score *Q* for a threshold t is given by,

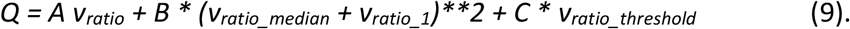

where A, B and C are user-definable parameters with defaults set by limited experimentation using a few test cases. (These default values are used throughout this work.) Values of the threshold *t* are tested and the value that maximizes the total score is used.

For iterations of segmentation after the first, the same approach is taken except that the density in all the regions that were above previous thresholds are set to zero and (by default) the estimate of remaining molecular volume is 20% of the volume in the previous iteration. This fraction of remaining volume is also user-adjustable and could likely be optimized as well.

Once a threshold is chosen and the resulting set of regions of connected density above that threshold are found, these regions are assembled into groups with members related by symmetry (if any is present). For a given region, possible symmetry-related regions are those for which randomly-placed points in the one region are mapped into the other region by a symmetry operation. Potential symmetry-related regions are sorted based on the fraction of randomly-placed points that mapped into the other region (with a minimum of 10% of these points mapping into the other region), and up to one symmetry-related region per symmetry operator is chosen to be a member of the symmetry-related group.

A unique set of density regions is chosen by picking one region from each symmetry-related group. The choice of regions is optimized to yield a compact structure and high connectivity. The compactness of the structure is represented by the radius of gyration of randomly-sampled points from all chosen regions. The connectivity of a set of regions is calculated based on finding the rms of the maximum gaps that would have to be spanned to connect each region to one central region, going through any number of regions in between. For any pair of regions, the gap is defined as the smallest distance between the randomly-sampled points in the two regions. For a set of regions, the overall gap is the largest of the individual gaps that would have to be crossed to go from one region to another, going through any other regions in the process. The overall connectivity score is the rms of these gaps for connections between each region and one central region.

The identity of the central region and all the other regions are adjusted to minimize both the connectivity score and the radius of gyration. The relative weighting of the two scores is determined by a user-definable parameter with a default value of weight on the radius of gyration of *weight_rad_gyr=0.1*. The weight on the radius of gyration is then normalized to the size of the molecule by multiplying this parameter by the largest cell dimension divided by 300 Å. (The salient feature is that the radius of gyration scales with the size of the molecule, the value of 300 Å is an arbitrary dimension).

The goal of the segmentation procedure described so far is to yield density corresponding to a single molecular unit. Such density could then be used as the basis for interpretation of one molecular unit in the map, and if the density covers a complete molecular unit then all the chains in that molecule would be completely represented and potentially fully traced. If the segmentation procedure yields only density corresponding to parts of several molecular units then complete chain tracing would not be possible. In this case it may be helpful to add regions of density around the initial segmented density in order to contain at least one full molecular unit. By default the *add_neighbors* procedure adds each region neighboring the initial regions that would lead to an increase in the overall radius of gyration of 1 Å or less.

Once the unique segmented density is identified, it is extracted in several ways. First, each individual region that is part of the unique segmented density is extracted and a new map is created that is slightly larger than the individual region and that contains this density surrounded by zero density. Second, the entire unique region is extracted, surrounded by zero density, and written out as a new map.

### Chain types to be examined

The chain types (protein, RNA, DNA) to be tested in model-building were automatically deduced from the contents of the sequence file using the *Phenix* method *guess_chain_types_from_sequences*. The one exception was that as the sequence for one structure (PDB entry 5Lij) which was built as a poly-alanine model, consisted entirely of the letter “A”, the chain type to be tested for this structure could not be assigned unambiguously and the keyword “chain_type=PROTEIN” was added when analyzing this reconstruction instead of the default of “chain_type=None”.

### Protein model-building

In our new core method for protein model-building, the *Phenix trace_chain* algorithm^14^ is used to build a polypeptide backbone through a map following high contour levels. These preliminary models are then improved by automatic iterative identification of secondary structure (see below) and refinement of the models including hydrogen-bond restraints representing this secondary structure with the *phenix.real_space_refine* approach^15^. As the connectivity in a map can sometimes be more evident at lower resolution, a series of maps blurred with different values of a blurring exponential factor *B*_*blur*_ ranging from 0 to 105 Å2 are created and each one is used in chain tracing.

Automatic identification of secondary structure is carried out by a feature-based method that is relatively insensitive to large errors so that even relatively poor models can be analyzed. Helices and sheets are found separately in our approach. Helices are identified from a helical geometry of C_α_ positions. Segments with 6 or more residues in length are considered, and C_α,i_ → C_α,i+ 3_ and C_α,i_ → C_α,i+ 4_ vectors are calculated. The helical rise from a C_α_ atom is taken to be the mean value of the length of the corresponding C_α,i_ → C_α,i+ 3_ and C_α,i_ → C_α,i+ 4_ vectors, divided by the mean number of intervening residues (3.5). The overall helical rise is taken to be the average rise over all suitable C_α_ atoms and the segment is rejected if the helical rise is not within 0.5 Å of the target of 1.54 Å. Then the mean of all C_α,i_ → C_α,i+ 3_ vectors is computed and the segment is considered helical if the mean dot product of individual vectors with the overall mean vector is at least 0.9 and no individual dot product is less than 0.3 (throughout this work, parameters are user-adjustable but values specified are used for all the work described here).

β-sheets are found by identification of parallel or antiparallel extended structure with at least 4 residues in each segment. For suitable C_α_ atoms within a single segment,C_α,i_ → C_α,i+ 3_ vectors are calculated. If the mean length of these vectors is within 1.5 Å of the target of 10 Å, the mean dot product of individual vectors with the overall average is at least 0.75 and no individual dot product is less than 0.5, the segment is considered as a possible strand. Two strands are considered part of the same sheet if the C_α_ in the strands can be paired 1:1 with rms side-to-side C_α,i_ →C_α,i’_ distances of 6 Å or less. As above, all parameters are user-adjustable.

Once helices, parallel or antiparallel strands are identified, the corresponding hydrogen-bonding pattern is used to generate restraints that are then used in real-space refinement.

A second method for model-building of proteins, previously used for crystallographic model-building, is based on examination of a map, typically at lower resolution, to identify features diagnostic of specific types of secondary structure. For example, at resolutions of 6 Å -8 Å, helices appear as distinct tubes of density^12^, and at resolutions near 4 Å, protein beta strands appear as approximately parallel tubes of density separated by a constant distance. These methods are implemented in the tool *phenix.find_helices_strands*^12^.

A third method used to build protein models into segmented regions was the *RESOLVE* model-building algorithm^13^ as implemented in *phenix.resolve*. This model-building method uses template matching to identify secondary structure elements (which can be protein helices or strands). Once secondary structure elements are identified, they are extended with tripeptide (or trinucleotide) fragments to create a full model.

Sequence assignment (matching of the sequence to residues in the model) for protein model-building is carried out using the tool *phenix.assign_sequence* which has been described recently^31^.

### Nucleic acid model-building

We have developed two approaches for nucleic acid model-building that have not previously been reported. The first is related to the template-matching methods for protein model-building described above^13^. DNA and RNA model-building are very similar and RNA model-building is described here. Regular A-form RNA helices (or B-form DNA helices) are identified with a convolution-based 6-dimensional search of a density map using regular base-paired templates 4 bases in length. Longer single-strands from base-paired templates of up to 19 bases are then superimposed on the templates that have been identified and portions that best match the density map are kept. These helical fragments are extended using libraries of trinucleotides based on 749 nucleotides in 6 RNA structures determined at resolutions of 2.5 Å or better (PDB entries 1gid, 1hr2, 3d2g, 4p95, 4pqv and 4y1j, using only one chain in each case) and filtered to retain only nucleotides where with the average B-value (atomic displacement parameter) was 50 Å^2^ or lower. Extension in the 5' direction using trinucleotides was done by superimposing the C4', C3', and C2' atoms of the 3' nucleotide base of the trinucleotide to be tested on the corresponding atoms of the 5' nucleotide in a placed segment, examining the map-model correlation for each trinucleotide, and choosing the one that best matches the map. A corresponding procedure was used for extension in the 3' direction. The matching of nucleic acid sequence to nucleotides in the model was carried with an algorithm similar to the one used in the tool *phenix.resolve* for protein sequence assignment^32^. Four to six conformations of each the bases were identified from the six RNA structures described above. For each conformation of each base, average density calculated at a resolution of 3 Å from the examples in these structures (after superimposing nucleotides using the C4', C3', and C2' atoms) was used as a density template for that base conformation. Then after a nucleotide chain was built with our algorithm, the map correlation between each of these templates and each position along the nucleotide chain is calculated and used to estimate the probability of each base at each position^32^. All possible alignments of the supplied sequence and each nucleotide chain built are considered and the best matches with a confidence of 95% or greater are considered matched. For these matched nucleotides the corresponding conformation of the matched base is then placed in the model. For residues where no match is found, the best-fitting nucleotide is used. For base-paired nucleotides, the same procedure is carried out, except that pairs of base-paired nucleotides are considered together, essentially doubling the amount of density information available for sequence identification.

A second new method for nucleic acid model-building used here is to build duplex RNA helices directly into the density map with the tool *phenix.build_rna_helices*. The motivation for this algorithm was that the nucleic-acid model-building approach described above, which builds the two chains of duplexes separately, frequently resulted in poorly base-paired strands. To build RNA helices directly, very similar overall strategies were used, except that the templates were all base-paired and base-paired nucleotides were always considered as a single unit. This automatically led to the same favorable base-pairing found in the structures used to derive the templates. The atoms used to superimpose chains and bases were the O4' and C3' atoms of one nucleotide and the C1' atom of the base-paired nucleotide. As in the previous method for sequence assignment, both bases in each base-paired set were considered together, leading again to a substantial increase in map-based information about the identities of bases in the model.

### Combining model information from different sources and removal of overlapping fragments

Model-building into local regions of density and into the entire asymmetric unit of the map normally yielded a set of partially overlapping segments. These segments were refined based on the sharpened map with the Phenix tool *phenix.real_space_refine*^15^. The refined segments were scored based on map-model correlation multiplied by the square root of the number of atoms in the segment (related to fragment scoring in *RESOLVE* model-building except that density at the coordinates of atoms was used instead of map-model correlation in that work^13^). Then a composite model was created from these fragments, starting from the highest-scoring one and working down, and including only non-overlapping parts of each new fragment considered, as implemented in the *Phenix tool phenix.combine_models*. When symmetry was used in reconstruction, all the symmetry-related copies of each fragment were considered in evaluating whether a particular part of a new fragment would overlap with the existing composite model.

### Construction and refinement of full model including reconstruction symmetry

In cases where symmetry had been used during the reconstruction process, we assumed that this symmetry was nearly perfect and applied this symmetry to the model that we generated. We began with our model that represented the asymmetric unit of the reconstruction. Then reconstruction symmetry was applied to this model and the model was refined against the sharpened map with the Phenix tool *phenix.real_space_refine*^15^. Finally one asymmetric unit of this final model was extracted to represent the unique part of the molecule and both the entire molecule with symmetry and the unique part are written out.

### Evaluation of model similarity to deposited structures

We developed the *Phenix tool phenix.chain_comparison* as a way of comparing the overall backbone (C_α_ or P atoms only) similarity of two models. The unique feature of this tool is that it considers each segment of each model separately so that it does not matter whether the chain is complete or broken into segments. Additionally the tool can separately identify segments that have matching C_α_ or P atoms that proceed in the same direction and those that are reversed as well as those that have insertions and deletions, as is common in low-resolution model-building. The *phenix.chain_comparison* tool also identifies whether the sequences of the two models match by counting the number of matching C_α_ or P atoms that are associated with matching residue names. These analyses are carried out with a default criteria that C_α_ or P atoms that are within 3 Å are matching and those further apart are not. This distance is arbitrary but was chosen to allow atoms to match in chains that superimpose secondary structure elements such as helices even if the register of the secondary structure elements do not superimpose exactly.

When comparing models corresponding to a reconstruction that has internal symmetry, the appropriate pairs of matching atoms may require application of that symmetry. Beginning with version 1.13-3015 of *Phenix*^28^, which we used for comparing models, the *phenix.chain_comparison* tool allows the inclusion of symmetry in the analysis.

### Datasets used

We selected reconstructions to analyze based on:

1. availability of a reconstruction in the EMDB as of Nov, 2017
2. resolution of the reconstruction 4.5 Å or better
3. presence of a unique deposited model in the PDB matching the reconstruction
4. consistent resolution in the PDB and EMDB
5. ability to use *Phenix* tools to automatically extract model and map from PDB and EMDB, apply symmetry if present in the metadata, and write the model

This resulted in 502 map-model pairs extracted from a total of 882 single-particle and helical reconstructions in the EMDB in this resolution range. (Note that only 660 of the 882 have one or more associated PDB entries).

After our initial analysis we further excluded reconstructions that had the following characteristics:

1. map-model correlation for the deposited map and deposited model of less than 0.3 after extraction of map and model and analysis with *phenix.map_model_cc* (18 reconstructions)
2. deposited model in the PDB represents less than half of the structure (9reconstructions)

This yielded the 476 map-model pairs that are described in this work. We downloaded the maps from the EMDB^16^ and used them directly in *phenix.map_to_model*, with the exception of one map (EMDB entry 6351). For EMDB entry 6351, a pseudo-helical reconstruction^33^, we could only deduce reconstruction symmetry for the part of the map corresponding to deposited model (PDB entry 3jar), so we used the *phenix.map_box* tool to cut out a box of density around the region defined by the deposited model, analyzed this map, and at the conclusion of the process translated the automatically-generated models to match the deposited map.

### Parameters used when running *phenix.map_to_model*

All of the reconstructions selected were analyzed with official version 1.13-3015 of *Phenix*, with all default values of parameters except those specifying file names for the reconstructed map, sequence file, and symmetry information, the resolution of the reconstruction, and any control parameters specific to the computing system and processing approach (e.g., the number of processors to use, queue commands,parameters specifying what parts of the calculation to carry out or what parts to combine in a particular job, and level of verbosity in output).

The *phenix.map_to_model* tool allows an analysis to be broken up into smaller tasks, followed by combining all the results to produce essentially the same result as would be obtained by running the entire procedure in one step. We used this approach for two purposes in this work. First, some of the datasets we analyzed required a very large amount of computer memory in certain stages of the analysis, and in particular in the map segmentation and final model construction steps. For other datasets such as ribosomes, the analysis could be substantially sped up by running smaller tasks on many processors.

## Computation

We carried out most of the analyses in this work on the *grizzly* high performance computing cluster at Los Alamos National Laboratory, typically with one task per node. Some of the analyses were carried out using a dedicated computing cluster at Lawrence Berkeley National Laboratory and some large analyses in particular were carried out on a machine with 1 TB of memory.

We monitored the CPU use for 175 of our analyses (generally smaller structures) that were each carried out in a single step on a single machine using a single processor. These analyses took from 15 minutes to 12 hours to complete on the *grizzly* cluster. For example, the analysis of EMDB entry 6630 in Fig. 2C required 3 CPU hours. We also monitored the CPU use for one of the largest structures (EMDB entry 9565), which required 129 CPU hours to complete.

## Code availability

The *phenix.map_to_model* tool and all other *Phenix* software is available along with full documentation in source and binary forms from the *Phenix* website^34^ as part of *Phenix* software suite^28^.

## Data availability

All the parameters, including resolution and specifications of reconstruction symmetry used in this work, are available at http://phenix-online.org/phenix_data/terwilliger/map_to_model_2018 along with all of the models produced.

## Acknowledgements

This work was supported by the NIH (grant GM063210 to PDA and TT) and the *Phenix* Industrial Consortium. This work was supported in part by the US Department of Energy under Contract No. DE-AC02-05CH11231 at Lawrence Berkeley National Laboratory. This research used resources provided by the Los Alamos National Laboratory Institutional Computing Program, which is supported by the U.S. Department of Energy National Nuclear Security Administration under Contract No. DE-AC52-06NA25396.

## Author contributions

PVA and OVS developed core tools for map segmentation and secondary structure restraints, PDA and PVA developed the real space refinement tools, TT developed the *map_to_model* tool and carried out the analyses, and PDA and TCT supervised the work.

## References

1. Kuhlbrandt, W. (2014). Science 343, 1443–1444.

2. Henderson, R. (2015) Archives of Biochemistry and Biophysics 58119–24.

3. Trabuco, L. G., Villa, E., Mitra, K., Frank, J., & Schulten, K. (2008). Structure 16 673–683.

4. Emsley, P., Lohkamp, B., Scott, W. G. & Cowtan, K. (2010). Features and development of Coot. Acta Cryst. D66 486–501.

5. Wang, R.Y., Kudryashev, M., Li, X, Egelman, E.H., Basler, M., Cheng, Y., Baker, D., DiMaio, F. (2015) Nature Methods 12 335–338.

6. Frenz, B., Walls., A.C., Egelman, E.H., Veesler, D., DiMaio, F. (2017). Nature Methods 14 797.

7. Chen M., Baldwin P.R., Ludtke, S.J., Baker M.L. (2015). J Struct Biol. 196 289–298.

8. DiMaio, F., Chiu, W. (2016). Methods. Enzymol. 679 255–276.

9. Zhou, N., Wang, H., Wang, J. (2017) Scientific Reports 7:2664.

10. Lawson, C. (2017). http://challenges.emdatabank.org/

11. Terwilliger T.C., Sobolev, O., Afonine P.V., Adams P.D. (2018). bioRxiv doi: https://doi.org/10.1101/247049.

12. Terwilliger, T. C. (2010). Acta Cryst. D66 268–275.

13. Terwilliger, T. C. (2003). Acta Cryst. D59 38–44.

14. Terwilliger, T. C. (2010). Acta Cryst. D66 285–294.

15. Afonine, P.V. et al. (2018). bioRxiv doi: https://doi.org/10.1101/249607.

16. Lawson, C.L., et al., (2016) Nucleic Acids Res. 44 D396–D403.

17. Berman H.M., et al., (2000). Nucleic Acids Research, 28 235–242.

18. Merk, A., et al., (2016). Cell 165: 1698–1707.

19. Borgnia, M.J. et al., (2016). Mol.Pharmacol. 89 645–651.

20. Pettersen, E.F., Goddard T.D., Huang C.C., Couch G.S., Greenblatt D.M., Meng E.C., Ferrin T.E. (2004). J. Comput. Chem. 25 1605–1612.

21. Li, Z. et al., (2017). Protein Cell, doi:10.1007/s13238-017-0456-9.

1. Schoebel, S. (2017). Nature 548 352–355.

23. Terwilliger, T.C., et al., (2007) Acta Cryst. D63 597–610.

24. Hryc, C.F., et al. (2017). Proc. Natl. Acad. Sci. USA 114 3103–3108.

25. Terwilliger T.C., Bricogne, G. (2014). Acta Cryst. D70 2533–2543.

26. Joosten, R.P., Joosten, K., Murshudov, G.N., Perrakis, A. (2012). Acta Cryst. D68:484–496.

27. Grabowski, M., et al. (2016). Acta Cryst. D72 1181–1193.

28. Adams P.D., et al. (2010). Acta Cryst. D66 213–221.

29. DeLaBarre, B., Brunger A.T. (2006). Acta Cryst. D62 92–32

30. Pintilie, G. D., Zhang, J., Goddard, T. D., Chiu, W., & Gossard, D. C., (2010). J. Struct. Biol. 170 427–438.

31. Terwilliger, T. C., et al. (2013). Acta Cryst. D69 2244–2250.

32. Terwilliger, T. C. (2003). Acta Cryst. D59 45–49.

33. Zhang, R., Alushin, G.M., Brown, A., Nogales, E. (2015). Cell 162 849–859.

34. Adams, P.D. (2017). http://www.phenix-online.org

